# Orai1- and Orai2-, but not Orai3-mediated I_CRAC_ is regulated by intracellular pH

**DOI:** 10.1101/2020.11.01.364299

**Authors:** Grigori Y. Rychkov, Fiona H. Zhou, Melissa K. Adams, Stuart M. Brierley, Linlin Ma, Greg J. Barritt

## Abstract

Three Orai (Orai1, Orai2 and Orai3) and two STIM (STIM1 and STIM2; <u>st</u>romal interaction <u>m</u>olecule) mammalian protein homologues constitute major components of the store-operated Ca^2+^ entry mechanism. When co-expressed with STIM1, Orai1, Orai2 and Orai3 form highly selective Ca^2+^ channels with properties of Ca^2+^ release activated Ca^2+^ (CRAC) channels. Despite the high level of homology between Orai proteins, CRAC channels formed by different Orai isoforms have distinctive properties, particularly with regards to Ca^2+^ dependent inactivation, inhibition/potentiation by 2-APB and sensitivity to reactive oxygen species. This study characterises and compares the regulation of Orai1, Orai2- and Orai3-mediated CRAC current (I_CRAC_) by intracellular pH. Using whole-cell patch clamping of HEK293T cells heterologously expressing Orai and STIM1 we show that I_CRAC_ formed by each Orai homologue has a unique sensitivity to changes in intracellular pH (pH_i_). Orai1-mediated I_CRAC_ exhibits a strong dependence on pH_i_ of both current amplitude and the kinetics of Ca^2+^ dependent inactivation. In contrast, Orai2 amplitude, but not kinetics, depends on pH_i_, whereas Orai3 shows no dependence on pH_i_ at all. Investigation of different Orai1-Orai3 chimeras suggests that pH_i_ dependence of Orai1 resides in both, the N-terminus and intracellular loop 2, and may also involve pH-dependent interactions with STIM1.

## INTRODUCTION

Store-operated Ca^2+^ channels, exemplified by Ca^2+^ release activated Ca^2+^ (CRAC) channel, provide an indispensable Ca^2+^ entry pathway governed by the filling state of the intracellular Ca^2+^ stores in most types of excitable and non-excitable cells (Putney, 1986; Prakriya *et al*., 2015). CRAC channels differ from all other known ion channels, as their molecular components the Orai and STIM (<u>st</u>romal interaction <u>m</u>olecule) proteins, which reside on the plasma membrane and the endoplasmic reticulum, respectively, have to interact in order to create functional channels in response to depletion of the intracellular Ca^2+^ stores (Prakriya *et al*., 2015). There are three Orai (Orai1, Orai2 and Orai3) and two STIM (STIM1 and STIM2) mammalian homologues, of which Orai1 and STIM1 are the best characterised (Liou *et al*., 2005; Roos *et al*., 2005; Feske *et al*., 2006; Vig *et al*., 2006). All Orai homologues form highly selective CRAC channels when co-expressed with STIM1 (Lis *et al*., 2007). The importance of physiological functions of Orai1 in cells of immune system, eccrine sweat glands, lacrimal glands, skeletal muscle and cells forming dental enamel have been deduced from studies of known human pathologies associated with mutations in *Orai1* gene and *Orai1*^*-/-*^ mice (Feske *et al*., 2006; Gwack *et al*., 2008; McCarl *et al*., 2010; Xing *et al*., 2014; Lacruz *et al*., 2015). Lack of known human pathologies associated with mutations in Orai2 and Orai3 has delayed clear identification of their physiological roles. However, there is evidence that Orai3 channels are overexpressed in some types of breast, lung and prostate cancers and are involved in the regulation of their growth and invasiveness (Motiani *et al*., 2010; Hoth *et al*., 2013; Motiani *et al*., 2013; Hasna *et al*., 2018; Azimi *et al*., 2019). Orai2 has been shown to modulate the magnitude of store-operated Ca^2+^ entry (SOCE) in T cells and mast cells thus playing a role in modulating T cell-mediated immune responses, mast cell degranulation and mast cell mediated anaphylaxis (Vaeth *et al*., 2017; Tsvilovskyy *et al*., 2018).

X-ray crystallography using *Drosophila* Orai revealed that the Ca^2+^ selective pore of CRAC channel is formed by a hexamer of Orai polypeptides (Hou *et al*., 2012). How many STIM1 polypeptides need to bind to Orai hexamer to form a functional channel, and exactly how they bind remains a point of contention (Nwokonko *et al*., 2017; Yen *et al*., 2019). Moreover, the number of STIM1 molecules bound to Orai1 in a functional channel is likely to vary, as inferred from patch clamping studies showing that different CRAC current (I_CRAC_) properties strongly depend on the STIM1:Orai1 expression ratios (Scrimgeour *et al*., 2009; Hoover *et al*., 2011). The most obvious I_CRAC_ characteristics affected by STIM1:Orai1 expression ratio, apart from the I_CRAC_ size, are fast Ca^2+^ dependent inactivation (FCDI) and Ca^2+^ dependent re-activation seen in the current traces recorded in response to voltage steps between -80 mV and -140 mV (Scrimgeour *et al*., 2009; Hoover *et al*., 2011; Rychkov, 2018). Our recent investigation identified a strong dependence of both FCDI and re-activation of Orai1/STIM1-mediated I_CRAC_ on intracellular pH (pH_i_), suggesting that pH_i_ affects Orai1-STIM1 interactions, effectively changing STIM1:Orai1 ratio in functional channels (Gavriliouk *et al*., 2017). Although the dependence of I_CRAC_ on pH, both extra- and intracellular, has previously been investigated in some detail, the exact mechanism of Orai1 dependence on pH_i_ remains poorly understood, and pH_i_ dependence of Orai2 and Orai3 has not been explored in a sufficient detail (Scrimgeour *et al*., 2012; Beck *et al*., 2014; Gavriliouk *et al*., 2017; Yu *et al*., 2018; Zhang *et al*., 2020).

In this study, we examined and compared pH_i_ dependence of Orai1-, Orai2- and Orai3-mediated I_CRAC_ using HEK293T cells heterologously expressing Orai and STIM1, whole-cell patch clamping, and bath applications of Na Propionate and NH_4_Cl to change intracellular pH. We found that Orai2 and Orai3 differ from Orai1 and from each other in their dependence on pH_i_. The amplitude of Orai2 current is affected by the changes in pH_i_ similarly to the amplitude of Orai1. However, unlike in Orai1, Orai2 FCDI remains unchanged at acidic pH_i_. In contrast to both Orai1 and Orai2, Orai3 is not sensitive to pH_i_ changes at all. Both Orai2 and Orai3 seem to lack Ca^2+^ dependent re-activation, which is prominent in Orai1 at acidic pH_i_ and low STIM1:Orai1 expression ratios (Scrimgeour *et al*., 2009; Gavriliouk *et al*., 2017). Mutagenesis of Orai1 suggests that the main difference between Orai1 and Orai3 with respect to their sensitivity to pH_i_ lies within N-terminus and intracellular loop 2 linking transmembrane domains 2 and 3, and possibly, in their interactions with STIM1.

## METHODS

### Cell culture and transfections

HEK293T human embryonic kidney cells (ATCC CRL 11268) were cultured at 37°C in 5% (v/v) CO_2_ in air in DMEM supplemented with 100 µM non-essential amino acids, 2 mM L-Glutamine and 10% fetal bovine serum. To express STIM1 and different Orai constructs cells plated on glass cover slips were transfected using Polyfect (Qiagen, Germany) transfection reagent according to the manufacturer’s instructions. All Orai constructs, except for Orai3 and Orai1-[Orai3-Nt-L2] chimera, were transfected at 4 STIM1 : 1 Orai ratio to obtain currents displaying clear FCDI (Scrimgeour *et al*., 2009). Our preliminary investigation suggested that FCDI of Orai3 is not affected by STIM1:Orai3 transfection ratio within 4:1 – 1:4 range (N. Scrimgeour, G. Rychkov; unpublished). However, to obtain the optimal size Orai3 currents it was found necessary to increase the amount of Orai3 plasmid in the transfection mixture up to 1 STIM1 : 1 Orai3 ratio. Therefore, Orai3 and Orai1-[Orai3-Nt-L2] chimera were co-transfected with STIM1 at 1:1 ratio. Patch clamp experiments were performed 24-48 hours following transfection.

### Mutagenesis

STIM1, Orai1, Orai2 and Orai3 were subcloned into pCMV-Sport6 and the GFP co-expressing vector pAdTrack-CMV as previously described (Scrimgeour *et al*., 2009). Orai1-[Orai3-Ct] (Orai1 with C-terminus of Orai3), Orai1-[Orai3-Nt] (Orai1 with N-terminus of Orai3) and Orai1-[Orai3-Nt-L2] (Orai1 with N-terminus (aa1-90) and intracellular loop2 (aa141-181) replaced with the corresponding segments of Orai3) chimeras and EE162/166QQ and H171Y mutations in Orai1were produced by PCR using Platinum Taq or Phusion High Fidelity DNA polymerase (Thermo Fisher Scientific (Life Technologies Australia), Vic, Australia). Orai1-CAD construct was produced in pCIneo vector by flanking WT Orail with STIM1 CAD (aa342-448) on its C-terminus after a linker of GGGGSGGGGS. All mutations were validated by DNA sequencing. For all details on mutagenesis please see Supporting Information. H155F mutant of Orai1 was generously provided by Dr L. Yue, University of Connecticut, USA. Orai1-[Orai3-L2] and Orai3-[Orai1-Nt-L2] chimeras and H134AOrai1 mutant were generously provided by Dr I. Frischauf and Prof C. Romanin, University of Linz, Austria.

### Electrophysiology

Whole-cell patch clamping was performed at room temperature using EPC-9 computer-based patch-clamp amplifier (HEKA Elektronik) and PatchMaster software (HEKA Elektronik). The control bath solution contained 140 mM NaCl, 4 mM CsCl, 10 mM CaCl_2_, 2 mM MgCl_2_, and 10 mM Na-Hepes adjusted to pH 7.4 with NaOH. To acidify intracellular pH, 15, 30 or 60 mM of NaCl in the control bath solution was replaced with 15, 30 or 60 mM Na Propionate correspondingly. To raise intracellular pH, 15 mM NaCl was replaced with 15 mM NH_4_Cl. Depletion of intracellular Ca^2+^ stores was achieved using 20 *μ*M Ins(3,4,5)P_3_ (Sigma) added to an internal solution containing, 135 mM caesium glutamate, 5 mM MgCl_2_, 1 mM MgATP, and 10 mM Na-HEPES adjusted to pH 7.2 with NaOH. Ca^2+^ in the pipette solution was buffered with either 10 mM EGTA or 10 mM BAPTA, as indicated. Patch pipettes were pulled from borosilicate glass and fire polished to give a pipette resistance between 2 and 4 MΩ. Series resistance did not exceed 12 MΩ and was 50-70% compensated. Following the establishment of whole cell configuration, the development of I_CRAC_ was monitored by applying voltage ramps from –120 to +120 mV every two seconds. All voltages shown are nominal voltages not corrected for the liquid junction potential between the bath and electrode solutions (approximately -18 mV when Na^+^ is the dominant external cation). The holding potential was 0 mV throughout for all constructs, except for Orai3 and Orai1-[Orai3-Nt-L2], which were recorded at a holding potential of 30 mV due to a shift of maximum open probability to more positive potentials. The EPC-9 amplifier compensated for cell capacitance automatically. Leakage was determined immediately before I_CRAC_ development using bath solution with 0 mM CaCl_2_ (10 mM MgCl_2_ was used to substitute for 10 mM CaCl_2_ in control bath solution) and subtracted using “leak subtraction” function of EPC9 amplifier.

### Data Analysis

Normalised instantaneous tail currents for voltage steps to -100 mV after 600 ms test pulses in the range of -120 to 80 mV were used to produce the apparent open probability (*P*_o_) curves. With the exception of the results shown in Fig. 6, for all apparent *P*_o_ curves the data were normalised to the amplitude of the tail current after a pulse between 40 and 80 mV. For Fig 6, the data were normalised to the amplitude of the tail current after a pulse to -120 mV. Where possible, data points were fitted with the Boltzmann distribution with an offset of the form:

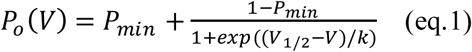

Where *P*_min_ is an offset, corresponding to the non-inactivating component of the current, *V* is the membrane potential, *V*_1/2_ is the half maximal activation potential (please note that *V*_1/2_ corresponds to the inflexion point of the *P*_o_ curve), and *k* is the slope factor. Otherwise, apparent *P*_o_ data points were fitted with a smooth curve using “cubic spline” procedure in GraphPad Prizm8 software (GraphPad Software, Inc., La Jolla, CA).

### Statistical Analysis

Data were analysed using GraphPad Prism 8 software. All values are reported as mean ± standard error (SEM). Statistical significance of differences between groups was determined using either unpaired Student’s t-test, One-way ANOVA or Two-way ANOVA as indicated.

## RESULTS

### Orai1-mediated I_CRAC_ is regulated by intracellular pH

All previous investigations of the dependence of I_CRAC_, either endogenous or mediated by ectopically expressed Orai1 and STIM1, on intracellular pH (pH_i_) were conducted by varying the pH of the patch pipette solution (Beck *et al*., 2014; Tsujikawa *et al*., 2015; Gavriliouk *et al*., 2017). A major disadvantage of this method is that intracellular solutions of different pH cannot be applied on the same cell. As a result, even relatively large changes in Orai/STIM current amplitude and/or kinetics only become obvious after a large number of experiments. To overcome this limitation, we used bath applications of different concentrations of Na Propionate (15, 30 and 60 mM) to reduce pH_i_ and NH_4_Cl (15 mM) to increase pH_i_ (Iino *et al*., 1994; Nitschke *et al*., 1996; Chen *et al*., 1998). First, we confirmed that the effects of extracellular application of Na Propionate and NH_4_Cl on Orai1/STIM1-mediated I_CRAC_ in HEK293T cells are comparable with previously published data on the effects of the acidic or alkaline intracellular solutions, respectively, with either EGTA or BATPA as the intracellular Ca^2+^ buffer in the pipette. With 10 mM EGTA in the pipette, application of Na Propionate to the bath resulted in a rapid and reversible inhibition of Orai1/STIM1 current (Fig. 1A, B, C). The current amplitude at -100 mV in the presence of 30 mM Na Propionate in the bath was 21.6±2.8% of the control (n=12; Fig. 1B), which was similar to the level of inhibition previously achieved using a pipette solution of pH 6.3 (Gavriliouk *et al*., 2017). In the presence of 15 mM NH_4_Cl in the bath, the amplitude of Orai1/STIM1 current increased 3.7±0.5 times (P<0.0001, One-way ANOVA) compared to the control (Fig. 1A, B, C). In contrast, when BAPTA was used to buffer Ca^2+^ in the pipette, NH_4_Cl had no effect on I_CRAC_ amplitude, and the reduction of the current amplitude in the presence of Na Propionate was significantly smaller, compared to the reduction seen with EGTA (Fig. 1D, E, F) (Fig. 1E, *cf*. Fig. 1B, P<0.001, Two-way ANOVA).

**Figure 1.**
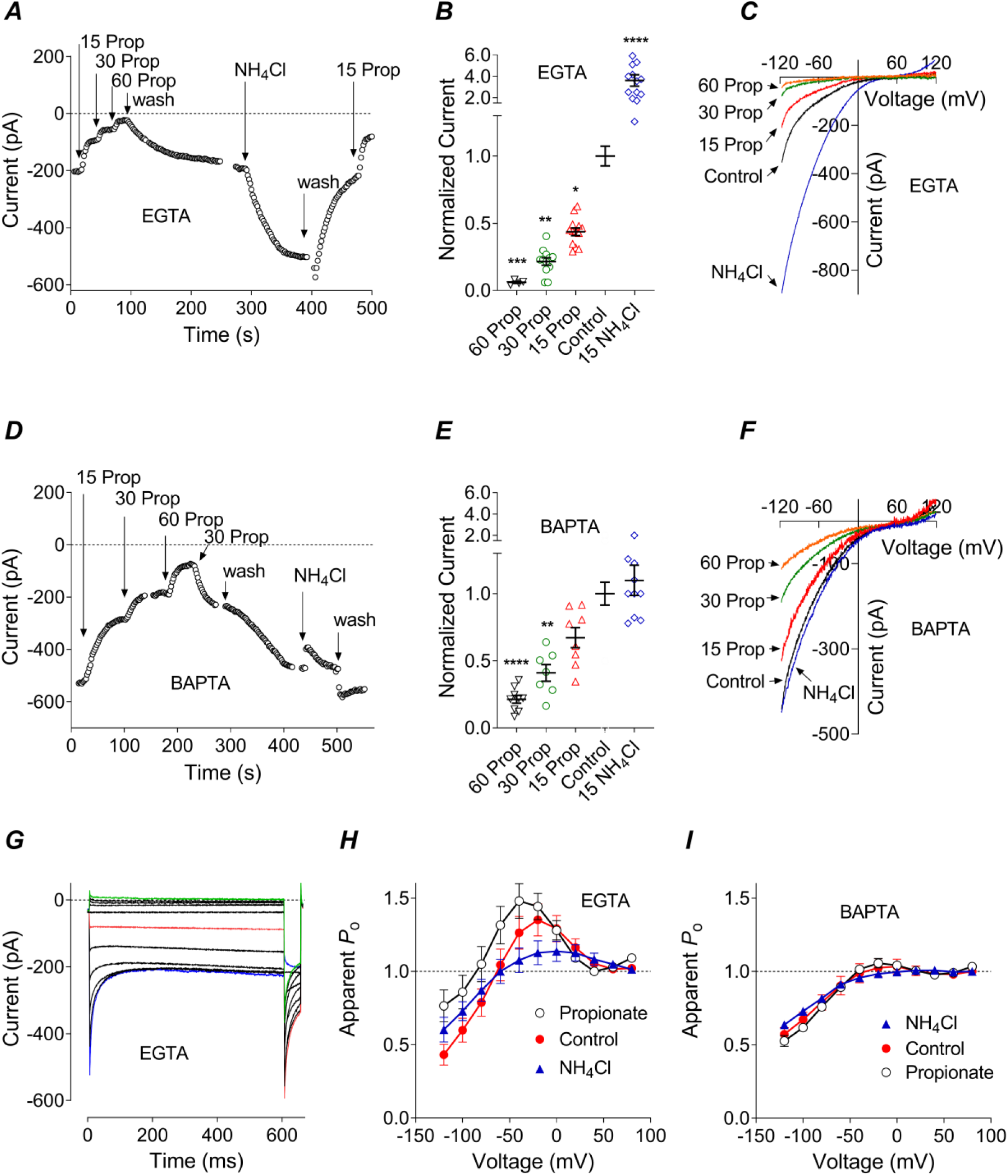
Investigation of Orai1 dependence on pH_i_ using Na Propionate and NH_4_Cl. A, D. Representative time course of Orai1-mediated I_CRAC_ amplitude during applications of Na Propionate and NH_4_Cl to the bath and either EGTA (A) or BAPTA (D) in the pipette solution. Each point represents I_CRAC_ amplitude at -100 mV, taken from I-V plots recorded in response to 100 ms voltage ramps from -120 to 120 mV. **B, E**. Average normalised amplitude of I_CRAC_ in the presence of Na Propionate or NH_4_Cl in the bath and either EGTA (B) or BAPTA (E) in the pipette solution. I_CRAC_ amplitude was measured at -100 mV as indicated in panel A and normalised to I_CRAC_ amplitude recorded in the control bath solution (One-way ANOVA, P<0.0001 (both B and E), with multiple comparisons to the control indicated within the panels). **C, F**. I_CRAC_ I-V plots recorded in response to 100 ms voltage ramps from -120 to 120 mV in the presence of Na Propionate or NH_4_Cl in the bath and either EGTA (C) or BAPTA (F) in the pipette solution. **G**. Representative I_CRAC_ traces showing tail currents at -100 mV following test pulses between -120 and 80 mV (see Experimental procedures). **H, I**. Apparent *P*_*o*_ curves generated using current traces similar to those shown in panel G, recorded in the control bath solution or in the presence of 15 mM Na Propionate or 15 mM NH_4_Cl in the bath and either EGTA (H) or BAPTA (I) in the pipette solution.

With EGTA in the pipette, applications of Na Propionate and NH_4_Cl not only affected the amplitude of I_CRAC_, but also its kinetics. To quantify I_CRAC_ fast Ca^2+^ dependent inactivation (FCDI) and re-activation we used normalised tail currents as previously described (Fig. 1G; see Experimental Procedures) (Scrimgeour *et al*., 2012; Gavriliouk *et al*., 2017). Intracellular acidification due to the presence of 15 mM Na Propionate in the bath caused an increase in I_CRAC_ re-activation, whereas intracellular alkalinisation by 15 mM NH_4_Cl reduced both, the FCDI and re-activation (Fig. 1H). As expected, intracellular BAPTA decreased FCDI and re-activation under control conditions (pH_i_ 7.3), and also reduced the effects of pH_i_ changes, induced by either Na Propionate or NH_4_Cl, on I_CRAC_ kinetics (Fig. 1I).

Overall, when EGTA was used to buffer Ca^2+^ in the pipette solution, changes in Orai1/STIM1-mediated I_CRAC_ amplitude and kinetics induced by Na Propionate and NH_4_Cl were comparable to the changes induced by acidic and alkaline intracellular solutions, correspondingly (Gavriliouk *et al*., 2017). Furthermore, if EGTA in the pipette solution was replaced with BAPTA, application of NH_4_Cl to the bath had no effect on the I_CRAC_ amplitude, and Na Propionate had little effect on I_CRAC_ kinetics (Fig. 1E, I), in good agreement with previous results obtained using pipette solutions with different pH (Gavriliouk *et al*., 2017).

### The amplitude but not FCDI of Orai2-mediated I_CRAC_ depends on pH_i_

Having established that bath applications of Na Propionate and NH_4_Cl accurately reproduce the effects of pH_i_ changes on I_CRAC_, we employed Na Propionate and NH_4_Cl to investigate the pH_i_ dependence of I_CRAC_ mediated by Orai2. The dependence of Orai2 current amplitude on pH_i_ with EGTA as Ca^2+^ buffer in the pipette was similar to that of Orai1 (Fig. 2A, B, C). When BAPTA was used to buffer intracellular Ca^2+^, the effect of intracellular acidification on Orai2 was somewhat weaker than on Orai1 (Fig. 2D, E, F) (Fig. 2E, *cf*. Fig. 1E, P=0.02, Two-way ANOVA). Nevertheless, 30 and 60 mM of Na Propionate in the bath solution induced a significant block of Orai2 current amplitude (Fig. 2D, E, F). HEK293T cells transfected with Orai2 and STIM1 at 1:4 ratio produced I_CRAC_ with properties different from those of Orai1-mediated I_CRAC_ in several respects. In agreement with previous investigations, the Orai2 currents exhibited slower kinetics of FCDI, compared to Orai1 (Fig. 2G) (Lis *et al*., 2007). The apparent *P*_o_ curve of Orai2 obtained under control conditions revealed a complete absence of I_CRAC_ re-activation, as the apparent *P*_o_ data could be fitted perfectly well with a single Boltzmann function (Fig. 2G, H; *cf*. Fig. 1I). Unlike Orai1, the kinetics of Orai2 current and the apparent *P*_o_ curve were unaffected by intracellular acidification (Fig. 2H). Intracellular alkalinisation by 15 mM NH_4_Cl reduced the slope of the apparent *P*_o_ curve and increased the non-inactivating component of the current (P=0.024, n=4, Student’s t-test) similarly to the replacement of EGTA in the pipette with BAPTA (Fig. 2H).

**Figure 2.**
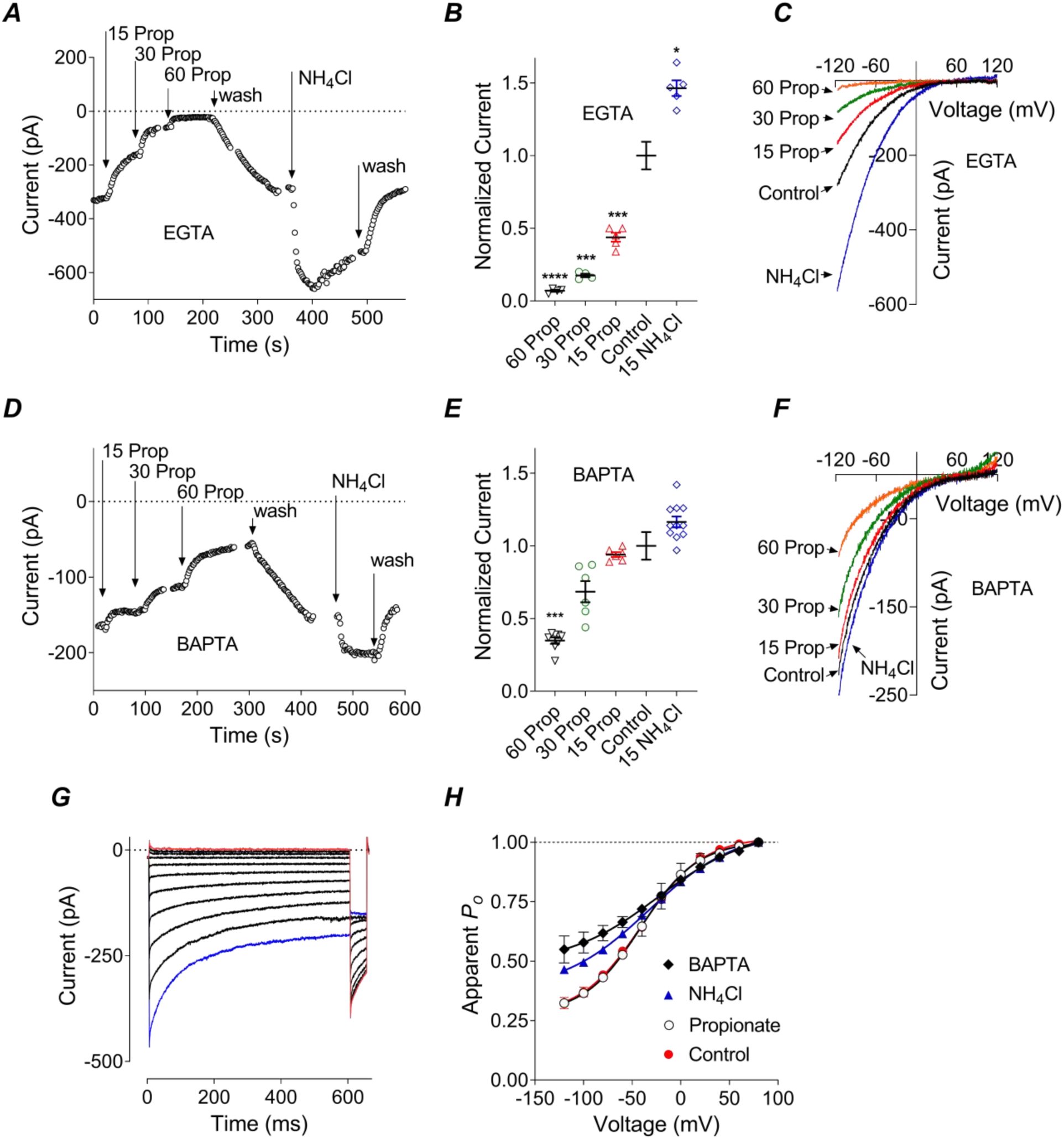
Dependence of Orai2-mediated I_CRAC_ on intracellular pH. A, D. Representative time course of Orai2-mediated I_CRAC_ amplitude during applications of Na Propionate and NH_4_Cl to the bath and either EGTA (A) or BAPTA (D) in the pipette solution. **B, E**. Average normalised amplitude of I_CRAC_ in the presence of Na Propionate or NH_4_Cl in the bath and either EGTA (B) or BAPTA (E) in the pipette solution (One-way ANOVA, P<0.001(B) and P<0.01 (E), with multiple comparisons to the control indicated within the panels). **C, F**. Orai2 I_CRAC_ I-V plots recorded in response to 100 ms voltage ramps from -120 to 120 mV in the presence of Na Propionate or NH_4_Cl in the bath and either EGTA (C) or BAPTA (F) in the pipette solution. **G**. Representative Orai2-mediated I_CRAC_ traces used to construct apparent *P*_*o*_ curves. **H**. Apparent *P*_*o*_ curves of Orai2 recorded in the control bath solution or in the presence of 15 mM Na Propionate or 15 mM NH_4_Cl in the bath and EGTA in the pipette solution, or in the control bath solution and BAPTA in the pipette, as indicated on the panel. Data points are fitted with Boltzmann function (eq 1, Experimental procedures).

### Orai3-mediated I_CRAC_ exhibits no dependence on pH_i_

Experiments similar to those described above but using HEK293T cells transfected with Orai3 and STIM1 revealed that I_CRAC_ mediated by Orai3 is largely independent of pH_i_ (Fig. 3). Orai3 current amplitude did not change appreciably in the presence of either Na Propionate or NH_4_Cl (Fig. 3A, B, C). If anything, intracellular acidification in the presence of BAPTA in the pipette caused an increase in I_CRAC_ amplitude (Fig. 3D, E, F). Similarly to Orai2, Orai3 did not exhibit any re-activation (see Experimental procedures; Fig. 3G, H). Furthermore, the Orai3 current kinetics and apparent *P*_o_ did not change in the presence of Na Propionate, but the non-inactivating component of the current (‘offset’ of the Boltzmann curve) increased in the presence of NH_4_Cl to 0.24±0.03, compared to 0.13±0.02 in control (Fig. 3H; P<0.01, Student’s t-test, n=7). As with Orai1 and Orai2, extracellular NH_4_Cl had an effect on Orai3 apparent *P*_o_ similar to that of intracellular BAPTA. However, the effects of either of them were less pronounced on Orai3 than on Orai1 or Orai2.

**Figure 3.**
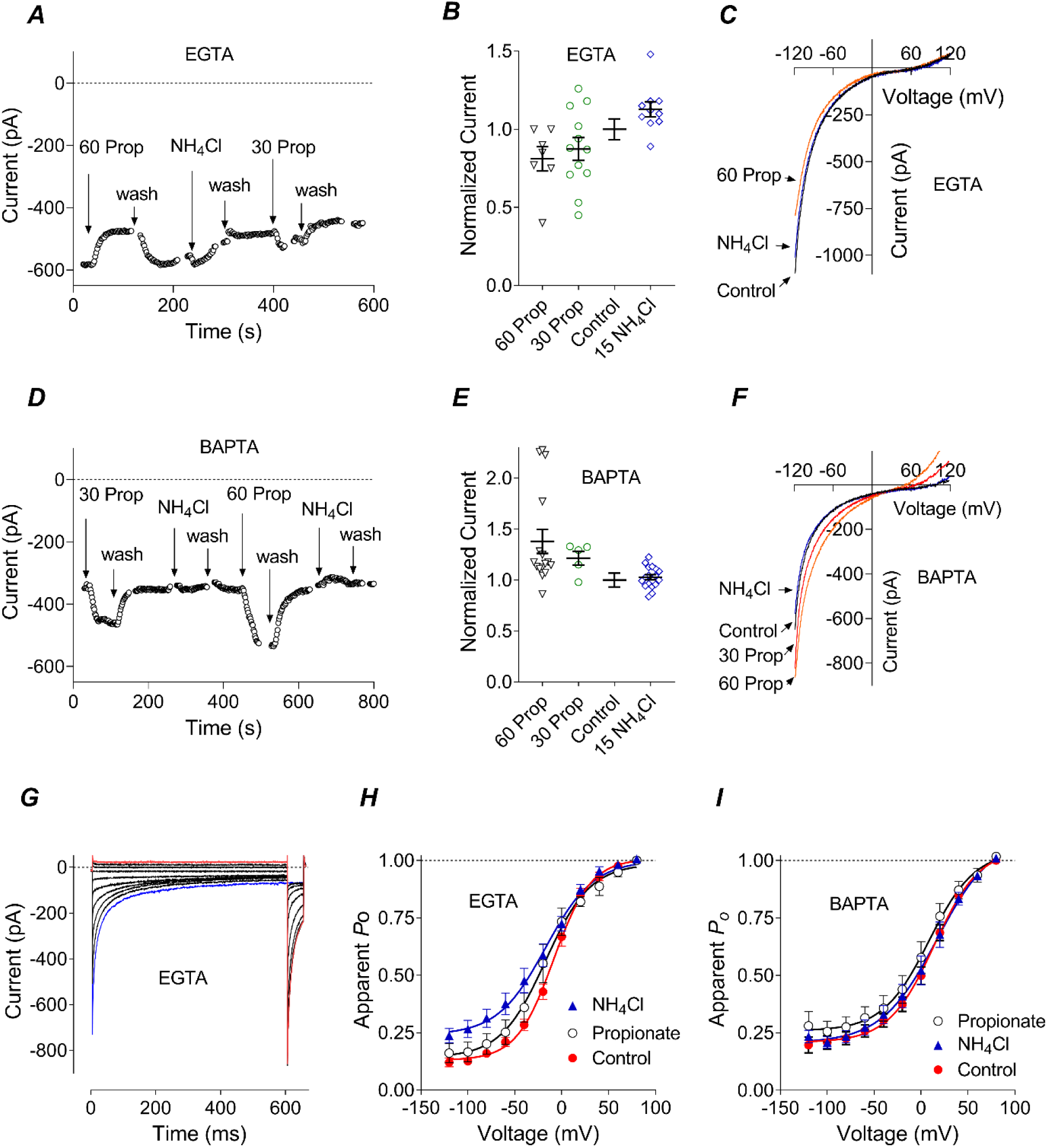
Orai3-mediated I_CRAC_ is largely independent of intracellular pH. A, D. Representative time course of Orai3-mediated I_CRAC_ amplitude during applications of Na Propionate and NH_4_Cl to the bath and either EGTA (A) or BAPTA (D) in the pipette solution. **B, E**. Average normalised amplitude of I_CRAC_ in the presence of Na Propionate or NH_4_Cl in the bath and either EGTA (B) or BAPTA (E) in the pipette solution (One-way ANOVA with multiple comparisons, no statistical difference between the control and any treatment (both, B and E)). **C, F**. Orai3 I_CRAC_ I-V plots recorded in response to 100 ms voltage ramps from - 120 to 120 mV in the presence of Na Propionate or NH_4_Cl in the bath and either EGTA (C) or BAPTA (F) in the pipette solution. **G**. Representative Orai3-mediated I_CRAC_ traces used to construct apparent *P*_o_ curves. **H, I**. Apparent *P*_o_ curves of Orai3 generated using current traces similar to those shown in panel G, recorded in the control bath solution or in the presence of 30 mM Na Propionate or 15 mM NH_4_Cl in the bath and either EGTA (H) or BAPTA (I) in the pipette solution. Data points are fitted with Boltzmann function (eq 1, Experimental procedures).

### Interaction between intracellular loop 2 and N-terminus plays a critical role in Orai1 dependence on pH_i_

Previous investigation of Orai1 dependence on pH_i_ by Tsujikawa *et al* suggested that histidine 155 in loop 2 mediates pH_i_ dependence of Na^+^ conductance through Orai1, determined in the absence of extracellular Ca^2+^ (Tsujikawa *et al*., 2015). Our investigation of H155F mutant of Orai1 (generously provided by Dr L. Yue, University of Connecticut) revealed that in the presence of Ca^2+^ in the bath solution, H155FOrai1 exhibits pH_i_ dependence similar to that of WT Orai1 (Fig. 4A-F). However, there were some differences between H155F Orai1 and WT Orai1 in the kinetics of FCDI and the effects of BAPTA. The FCDI of H155F mediated current was slower (Fig. 4G, *cf*. Fig. 1G; time constant of current inactivation at - 120 mV, τ, increased from 10.3±1.1 ms in WTOrai1 to 23.5±2.1 ms in H115FOrai1 mutant; P=0.0014, n=4, Student’s t-test) and the replacement of intracellular EGTA with BAPTA had a lesser effect on FCDI and re-activation of H155F, compared to WT Orai1 (Fig. 4I, *cf*. Fig. 1I). These results imply that in the presence of 10 mM Ca^2+^ in the bath solution H155F differs from WT Orai1 in its regulation by intracellular Ca^2+^, but not pH.

**Figure 4.**
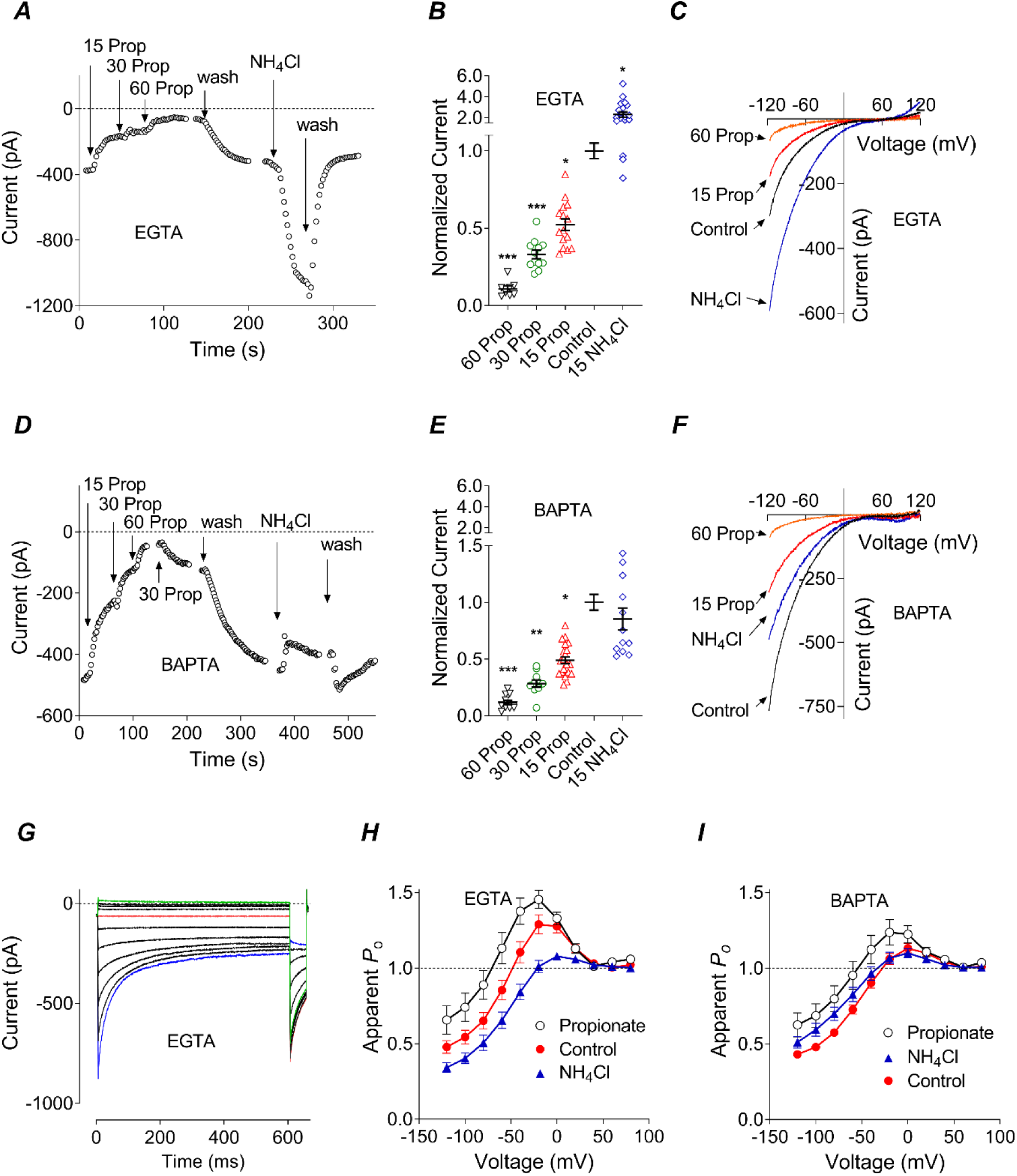
H115F mutation in the intracellular loop2 of Orai1 has no effect on the pH_i_ dependence Orai1-mediated I_CRAC_. **A, D**. Representative time course of H115F-Orai1-mediated I_CRAC_ amplitude during applications of Na Propionate and NH_4_Cl to the bath and either EGTA (A) or BAPTA (D) in the pipette solution. **B, E**. Average normalised amplitude of H115F-Orai1 I_CRAC_ in the presence of Na Propionate or NH_4_Cl in the bath and either EGTA (B) or BAPTA (E) in the pipette solution (One-way ANOVA, P<0.0001 (B) and P<0.001 (E), with multiple comparisons to the control indicated within the panels). **C, F**. H115F-Orai1 I_CRAC_ I-V plots recorded in response to 100 ms voltage ramps from -120 to 120 mV in the presence of Na Propionate or NH_4_Cl in the bath and either EGTA (C) or BAPTA (F) in the pipette solution. **G**. Representative H115F-Orai1 I_CRAC_ traces used to construct apparent *P*_*o*_ curves. **H, I**. Apparent *P*_o_ curves generated using current traces similar to those shown in panel G, recorded in the control bath solution or in the presence of 15 mM Na Propionate or 15 mM NH_4_Cl in the bath and either EGTA (H) or BAPTA (I) in the pipette solution.

A logical assumption from the lack of Orai3 dependence on pH_i_ is that some structural differences in the intracellular domains, or transmembrane domains of Orai1 and Orai3 internally accessible to protons are responsible for the stark difference between their dependence on pH_i_. The transmembrane domains of all Orai isoforms are highly conserved and all the difference between Orai1 and Orai3 in the potential protonation sites lies within N- and C-termini and the intracellular loop 2 (Deng *et al*., 2009; Frischauf *et al*., 2011). To determine which components are responsible for Orai1 dependence on pH_i_ we first separately replaced N- and C-termini of Orai1 with those of Orai3. Both chimeras, when co-expressed with STIM1 at 1:4 ratio, produced I_CRAC_ with pH_i_ dependence of Orai1 (Fig. S1). Orai1-[Orai3-Ct] construct (Orai1 with C-terminus of Orai3) exhibited FCDI and re-activation similar to that of Orai1 (Fig. S1A, *iii*). Orai1-[Orai3-Nt] chimera (Orai1 with N-terminus of Orai3), however, exhibited FCDI similar to that of Orai3, while retaining re-activation of Orai1 (Fig. S1B, *iii*).

Replacing Orai1 intracellular loop 2 with the intracellular loop 2 of Orai3 (Orai1-[Orai3-L2] chimera) produced I_CRAC_ with properties characteristic of Orai1 (Fig. S1C), which exhibited both FCDI and re-activation, and a clear dependence of the current amplitude on pH_i_ (Fig. S1C). However, compared to the WT Orai1, Orai1-[Orai3-L2] chimera was less sensitive to the changes in pH_i_, particularly when BAPTA was present in the pipette solution (Fig. S1C*ii, cf*. Fig 1C; P<0.0001, two-way ANOVA). This suggested that the intracellular loop 2 was contributing to the Orai1 regulation by pH_i_, but it was not clear whether any specific protonatable amino acids in loop 2 mediated this dependence. There are only two potentially protonatable amino acid residues in Orai1 loop 2, which are present in Orai2, but absent in Orai3 - glutamates 162 and 166 (Deng *et al*., 2009; Frischauf *et al*., 2011). In addition, there is one histidine in Orai1 loop 2 (H171), which is absent in both, Orai2 and Orai3 (Deng *et al*., 2009). However, mutating both glutamates simultaneously to glutamines (EE162/166QQ) or histidine to tyrosine (H171Y) in Orai1, to replace these protonatable residues with their equivalents in Orai3, had no effect on Orai1 dependence on pH_i_ (Fig. S2).

Previous search for the structures responsible for the differences between FCDI kinetics of Orai1 and Orai3 revealed that FCDI depends on the cooperative action of all intracellular domains, C- and N-termini and the intracellular loop 2, and the replacement of all 3 intracellular domains in Orai1 with the Orai3 counterparts produced I_CRAC_ exhibiting Orai3 FCDI (Frischauf *et al*., 2011). It is possible, that pH_i_ dependence of Orai1 also arises from cooperative action of 2 or 3 intracellular domains. Replacing both intracellular loop 2 and N-terminus in Orai1with those of Orai3 (Orai1-[Orai3-Nt-loop2] chimera) resulted in I_CRAC_ with properties resembling Orai3 (Fig. 5A-I). I_CRAC_ mediated by Orai1-[Orai3-Nt-loop2] chimera co-transfected with STIM1 exhibited pronounced FCDI, absence of re-activation and a complete lack of I_CRAC_ amplitude dependence on pH_i_ (Fig. 5A-I). This confirmed that Orai1 regulation by pH_i_ requires interaction between N-terminus and the intracellular loop 2.

**Figure 5.**
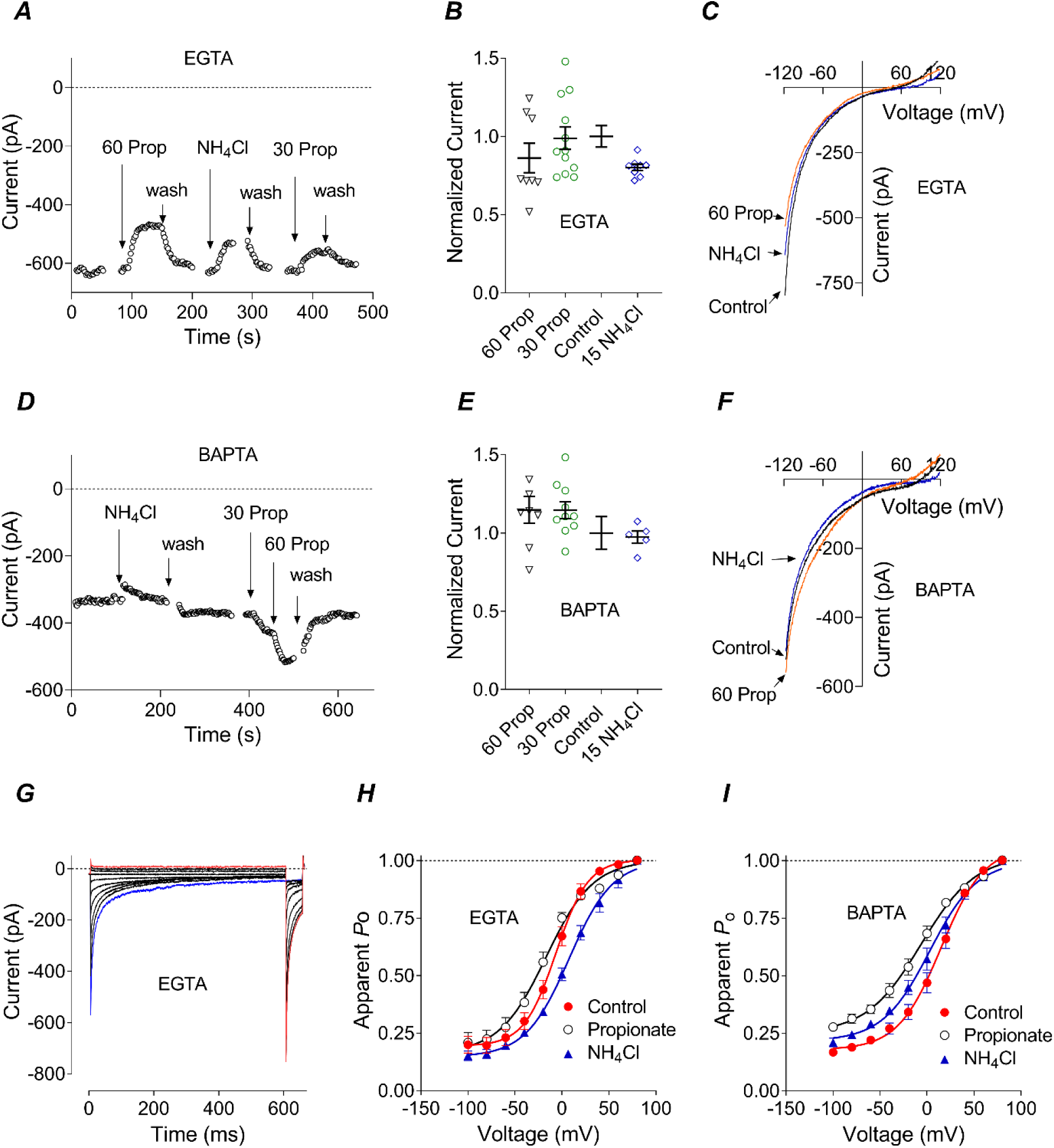
Replacement of Orai1 N-terminus and intracellular loop2 with N-terminus and loop2 of Orai3 renders Orai1-mediated I_CRAC_ independent of pH_i_. **A, D**. Representative time course of Orai1- [Orai3-Nt-L2] chimera I_CRAC_ amplitude during applications of Na Propionate and NH_4_Cl to the bath and either EGTA (A) or BAPTA (D) in the pipette solution. **B, E**. Average normalised amplitude of Orai1- [Orai3-Nt-L2] I_CRAC_ in the presence of Na Propionate or NH_4_Cl in the bath and either EGTA (B) or BAPTA (E) in the pipette solution (One-way ANOVA with multiple comparisons, no statistical difference between the control and any treatment (both, B and E)). **C, F**. Orai1-[Orai3-Nt-L2] I_CRAC_ I-V plots recorded in response to 100 ms voltage ramps from -120 to 120 mV in the presence of Na Propionate or NH_4_Cl in the bath and either EGTA (C) or BAPTA (F) in the pipette solution. **G**. Representative Orai1-[Orai3-Nt-L2] I_CRAC_ traces used to construct apparent *P*_*o*_ curves. **H, I**. Apparent *P*_o_ curves of Orai1-[Orai3-Nt-L2] chimera generated using current traces similar to those shown in panel G, recorded in the control bath solution or in the presence of 30 mM Na Propionate or 15 mM NH_4_Cl in the bath and either EGTA (H) or BAPTA (I) in the pipette solution. Data points are fitted with Boltzmann function (eq 1, Experimental procedures).

**Figure 6.**
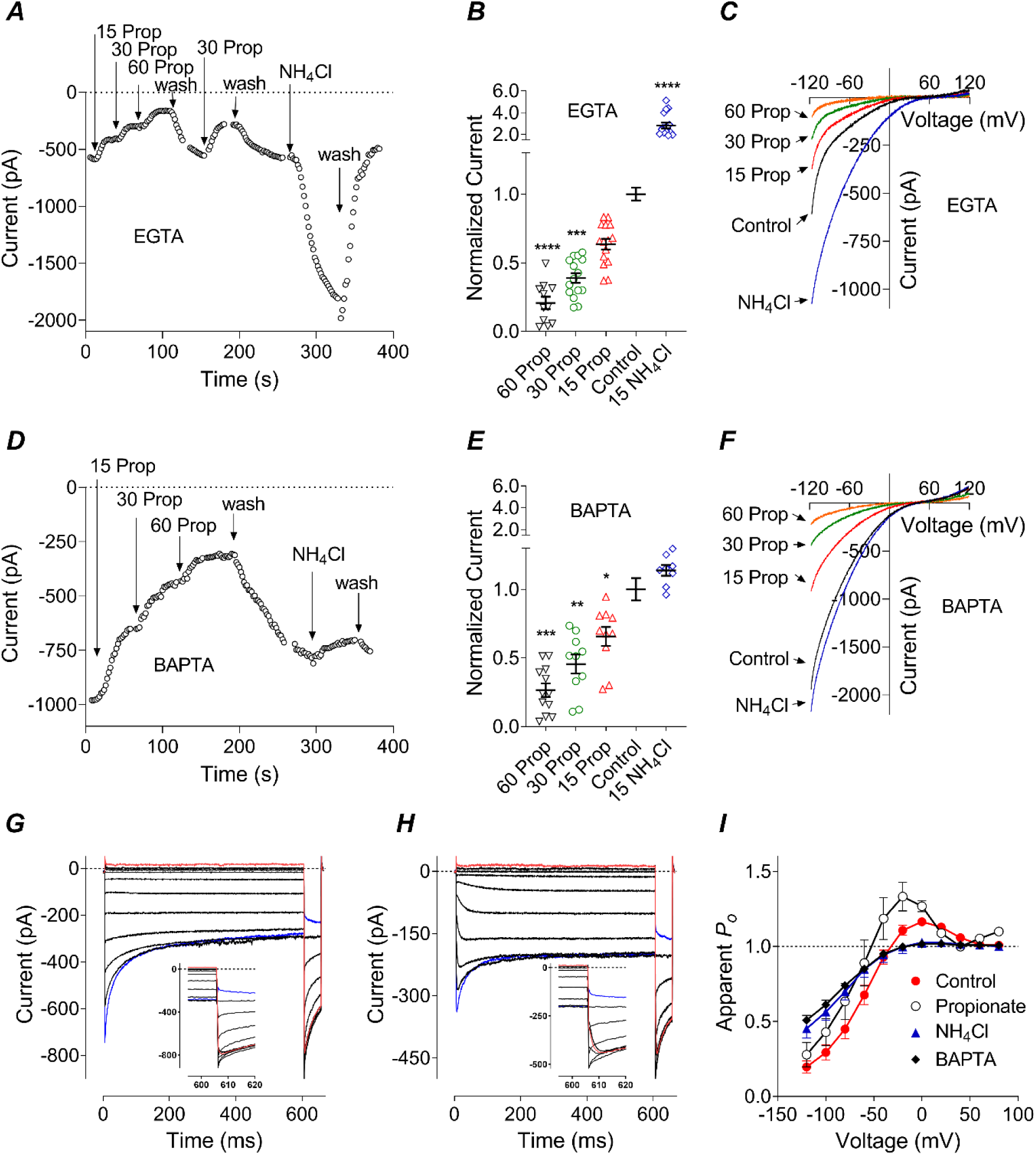
Replacement of Orai3 N-terminus and intracellular loop2 with N-terminus and loop2 of Orai1 affords pH_i_ dependence to Orai3-mediated I_CRAC_. **A, D**. Representative time course of Orai3- [Orai1-Nt-loop2] chimera I_CRAC_ amplitude during applications of Na Propionate and NH_4_Cl to the bath and either EGTA (A) or BAPTA (D) in the pipette solution. **B, E**. Average normalised amplitude of Orai3- [Orai1-Nt-L2] I_CRAC_ in the presence of Na Propionate or NH_4_Cl in the bath and either EGTA (B) or BAPTA (E) in the pipette solution (One-way ANOVA, P<0.0001 (both B and E), with multiple comparisons to the control indicated within the panels). **C, F**. Orai3-[Orai1-Nt-L2] I_CRAC_ I-V plots recorded in response to 100 ms voltage ramps from -120 to 120 mV in the presence of Na Propionate or NH_4_Cl in the bath and either EGTA (C) or BAPTA (F) in the pipette solution. **G, H**. Representative Orai3-[Orai1-Nt-L2] I_CRAC_ traces used to construct apparent *P*_*o*_ curves in the control bath solution (G) and bath solution with 15 mM propionate (H). The insets show peak tail currents at expanded time scale (also see Fig. S3) **I**. Apparent *P*_o_ curves of Orai3-[Orai1-Nt-L2] chimera generated using current traces similar to those shown in panel G, recorded in the control bath solution or in the presence of 15 mM Na Propionate or 15 mM NH_4_Cl in the bath and EGTA in the pipette solution or in the control bath solution and BAPTA in the pipette, as indicated on the panel.

To further elucidate the role of these intracellular domains, pH_i_ dependence was investigated using an Orai3 construct in which the N-terminus and the intracellular loop 2 of Orai3 were replaced with those of Orai1 (Orai3-[Orai1-Nt-loop2] chimera (generously provided by Dr I. Frischauf and Prof C. Romanin, University of Linz, Austria) (Frischauf *et al*., 2011). Application of Na Propionate or NH_4_Cl to the cells co-expressing Orai3-[Orai1-Nt-loop2] construct with STIM1 revealed that replacement of both, the N-terminus and the intracellular loop 2 of Orai3 with those of Orai1 was sufficient to render this chimera dependent on pH_i_ (Fig. 6). The results indicate that the extent of inhibition of I_CRAC_ mediated by Orai3-[Orai1-Nt-loop2] chimera induced by intracellular acidification, or the potentiation of I_CRAC_ mediated by this chimera due to alkalinisation were not statistically different from those observed in Orai1-mediated I_CRAC_ (Fig. 6A-F). Furthermore, the apparent *P*_o_ curves of Orai3-[Orai1-Nt-loop2]-mediated I_CRAC_ displayed dependence on pH_i_ similar to that of Orai1 (Fig. 6I, *cf*. Fig. 1H). There was, however, a significant difference in the kinetics of I_CRAC_ re-activation (Fig. 6G, H). Orai1 I_CRAC_ re-activation at negative potentials occurs on a time scale of hundreds of milliseconds (Scrimgeour *et al*., 2009). Compared to WT Orai1, re-activation of Orai3-[Orai1-Nt-loop2] I_CRAC_ was more than 100 times faster (τ∼2 ms in Orai3-[Orai1-Nt-loop2] *vs* τ∼ 360 ms in WT Orai1, at -100 mV) (Fig. 6G, H, insets and Fig. S3; *cf*. Fig 7G, H). Nevertheless, this fast re-activation was more pronounced at acidic pH_i_ (Fig. 6H, I; Fig. S3), similarly to the slow re-activation observed in Orai1 I_CRAC_ (Fig. 1H) (Gavriliouk *et al*., 2017).

**Figure 7.**
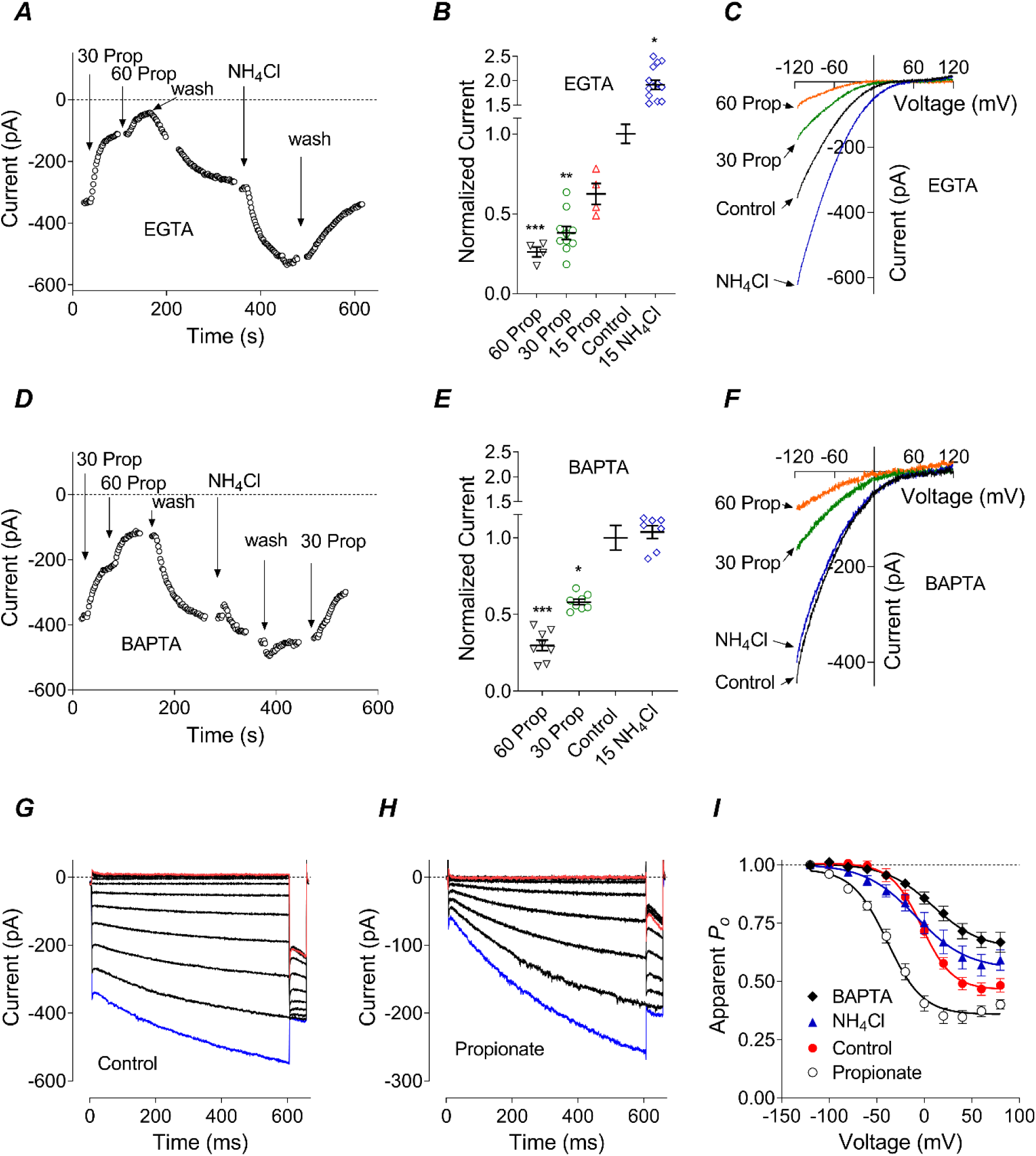
Orai1-CAD chimera largely retains pH_i_ dependence of WT Orai1-STIM1-mediated I_CRAC_. **A, D**. Representative time course of Orai1-CAD-mediated I_CRAC_ amplitude during applications of Na Propionate and NH_4_Cl to the bath and either EGTA (A) or BAPTA (D) in the pipette solution. **B, E**. Average normalised amplitude of Orai1-CAD I_CRAC_ in the presence of Na Propionate or NH_4_Cl in the bath and either EGTA (B) or BAPTA (E) in the pipette solution (One-way ANOVA, P<0.001 (B and E), with multiple comparisons to the control indicated within the panels). **C, F**. Orai1-CAD I_CRAC_ I-V plots recorded in response to 100 ms voltage ramps from -120 to 120 mV in the presence of Na Propionate or NH_4_Cl in the bath and either EGTA (C) or BAPTA (F) in the pipette solution. **G, H**. Representative Orai1-CAD I_CRAC_ traces used to construct apparent *P*_*o*_ curvesin the control bath solution (G) and bath solution with 15 mM propionate (H). **I**. Apparent *P*_o_ curves of Orai1-CAD recorded in the control bath solution, in the presence of 15 mM Na Propionate or 15 mM NH_4_Cl in the bath and EGTA in the pipette solution, or in the control bath solution and BAPTA in the pipette, as indicated on the panel. Data points are fitted with Boltzmann function (eq 1, Experimental procedures). Please note that the data on this panel were normalised to the amplitude of the tail current after a pulse to -120 mV, unlike all other *P*_o_ curves above where the data were normalised to the amplitude of the tail current after a pulse between 40 and 80 mV. Data points are fitted with Boltzmann function.

### STIM1 contributes to pH_i_ dependence of Orai1/STIM1-mediated I_CRAC_

Prominent Ca^2+^ dependent re-activation of Orai1/STIM1-mediated I_CRAC_ induced by intracellular acidification is prevalent in cells expressing high levels of Orai1 relative to STIM1 even at normal pH_i_, suggesting that pH_i_ affects Orai1-STIM1 interactions, effectively changing STIM1:Orai1 ratio in functional channels (Scrimgeour *et al*., 2009; Hoover *et al*., 2011; Gavriliouk *et al*., 2017). Both, the N-terminus and the intracellular loop 2 of Orai1 have been shown to contain sites interacting with STIM1 (for review see (Lunz *et al*., 2019)), and it is possible that protonation of the specific residues in Orai1 affects Orai1-STIM1 interactions. It is equally possible that some of the protonatable residues that mediate I_CRAC_ dependence on pH_i_ are localised in STIM1. Previous investigation of the effects of mutations neutralising negative charges in the STIM1 inactivation domain (ID_STIM_) revealed that ID_STIM_ is not involved in pH_i_ dependence of I_CRAC_ amplitude (Gavriliouk *et al*., 2017). To narrow down the regions of STIM1 that could contain relevant protonatable sites, we investigated the pH_i_ dependence of I_CRAC_ mediated by an Orai1-CAD chimera (Fig. 7) (CAD; <u>C</u>RAC <u>a</u>ctivation <u>d</u>omain - minimal STIM1 fragment (342-448aa) required for functional I_CRAC_ (Park *et al*., 2009)). This consisted of a full length Orai1 fused at the C-terminus with CAD. The amplitude of Orai1-CAD current showed strong dependence on pH_i_ for both intracellular Ca^2+^ buffers, EGTA (P<0.0001, One-way ANOVA) and BAPTA (P<0.001, One-way ANOVA) (Fig. 7A-F). However, this dependence was somewhat weaker than that of Oria1/STIM1-mediated I_CRAC_ (Fig. 7B, *cf*. Fig 1B; P<0.01, Two-way ANOVA). As expected, HEK293T cells transfected with Orai1-CAD cDNA produced a constitutively active I_CRAC_ which lacked FCDI (Fig. 7G)(McNally *et al*., 2012). Orai1-CAD mediated I_CRAC_ did, however, exhibit re-activation at negative potentials which was more prominent at acidic pH_i_ (Fig. 7G, H). Intracellular acidification promoted closure of the channel at positive potentials, shifting apparent *P*_o_ curve to hyperpolarising potentials and decreasing non-inactivating component of the current (Fig. 7G, H, I). Intracellular alkalinisation reduced the slope of the apparent *P*_o_ curve and increased the steady-state current, similarly to BAPTA (Fig. 7I). Increased steady-state current in the presence of BAPTA confirmed that I_CRAC_ re-activation is a Ca^2+^ dependent process, similarly to FCDI. The apparent *P*_o_ data could be fitted well with a single Boltzmann function with a maximum around -100 mV and minimum at about 60 mV (Fig. 7I).

Apparent *P*_o_ data in Fig. 7I were obtained by normalising instantaneous tail currents to the amplitude of the tail current corresponding to -120 mV, whereas the apparent *P*_o_ data for all other constructs presented in this paper were normalised to the tail currents obtained after voltage steps to 40-80 mV. The Orai1-CAD data shown in Fig. 7I but normalised to the tail current obtained after voltage step to 60 mV, is presented in Fig. S3. The curves in Fig. S3 suggests that the changes in I_CRAC_ kinetics and the apparent *P*_o_ curves induced by pH_i_ acidification are mostly due to pH_i_ dependence of re-activation, rather than FCDI (Fig. S4, *cf*. Fig. 1H).

The results above suggest that if STIM1 contains any protonatable amino acid residues relevant for I_CRAC_ dependence on pH_i_, they must be localised within CAD. Mutating CAD is problematic due to likely effects of mutations on CAD interaction with Orai1. Therefore, to help decipher a possible role of STIM1 in I_CRAC_ dependence on pH_i_, we used H134A Orai1 mutant (generously provided by Dr I. Frischauf and Prof C. Romanin, University of Linz, Austria) which does not require STIM1 to function (Zhou *et al*., 2016; Frischauf *et al*., 2017; Lunz *et al*., 2019). H134A Orai1 largely exhibits the properties of the WT Orai1 but is constitutively open and lacks FCDI or re-activation (Frischauf *et al*., 2017) (Fig. 8). Intracellular acidification induced by bath applications of 30 and 60 mM Na Propionate inhibited I_CRAC_ mediated by H134A Orai1 mutant expressed in HEK293T cells without STIM1 (Fig. 8A-F). However, this inhibition was significantly weaker than inhibition observed in WT Orai1 (Fig. 8, *cf*. Fig. 1; P<0.0001, two-way ANOVA). Furthermore, intracellular alkalinisation had no effect on the current amplitude, and there was no difference between pH_i_ dependence of H134AOrai1 currents recorded using either EGTA or BAPTA as intracellular Ca^2+^ buffer (Fig. 8B, E).

**Figure 8.**
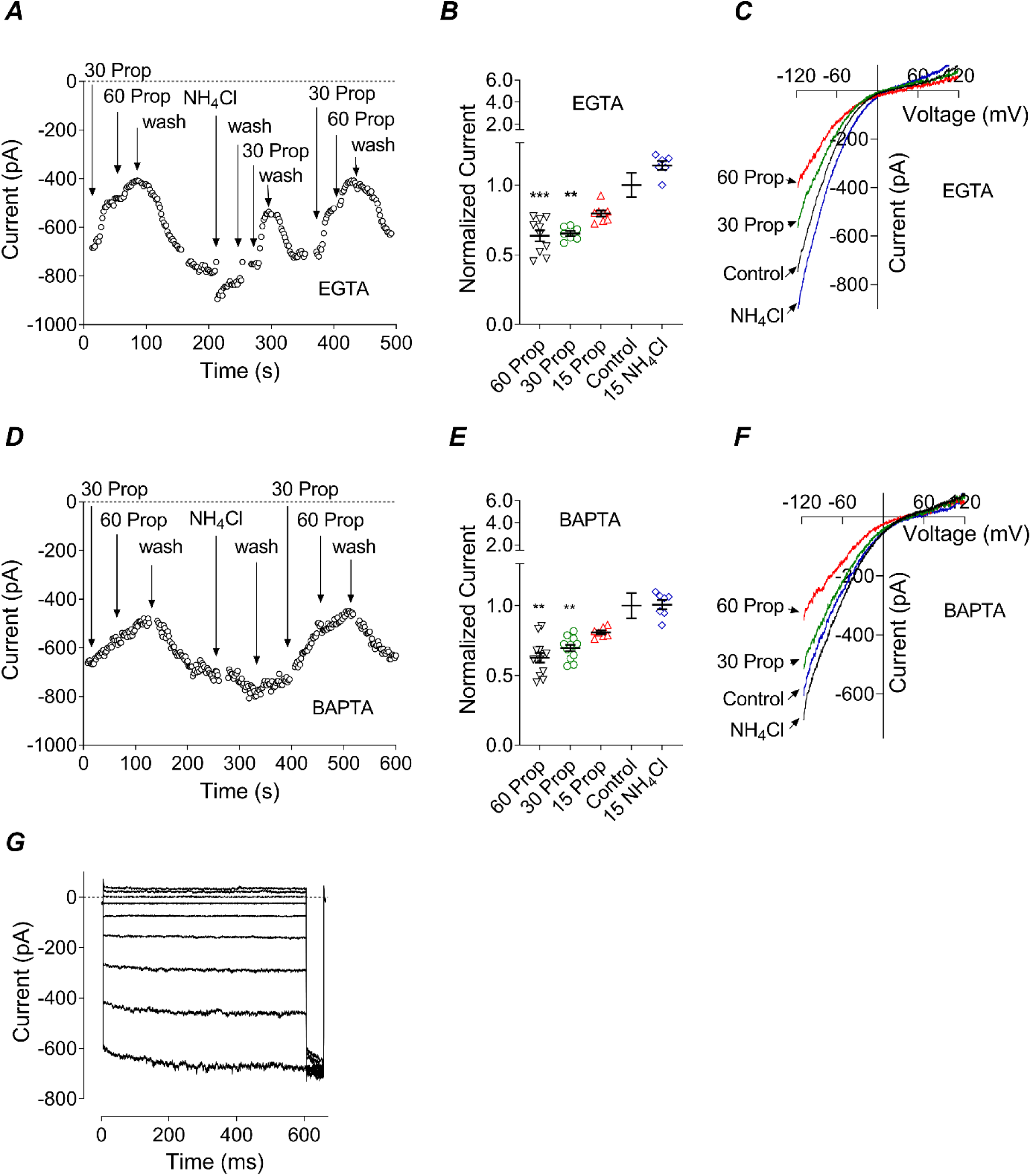
H134A mutation in Orai1 significantly attenuates pH_i_ dependence Orai1-mediated I_CRAC_. **A, D**. Representative time course of H134A-Orai1-mediated I_CRAC_ amplitude during applications of Na Propionate and NH_4_Cl to the bath and either EGTA (A) or BAPTA (D) in the pipette solution. **B, E**. Average normalised amplitude of H134A -Orai1 I_CRAC_ in the presence of Na Propionate or NH_4_Cl in the bath and either EGTA (B) or BAPTA (E) in the pipette solution (One-way ANOVA, P<0.001 (B and E), with multiple comparisons to the control indicated within the panels). **C, F**. H134A-Orai1 I_CRAC_ I-V plots recorded in response to 100 ms voltage ramps from -120 to 120 mV in the presence of Na Propionate or NH_4_Cl in the bath and either EGTA (C) or BAPTA (F) in the pipette solution. **G**. H134A-Orai1 I_CRAC_ recorded in response to the voltage protocol used to obtain the *P*_o_ curves for all other Orai constructs. It shows lack of both FCDI and re-activation.

## DISCUSSION

In this work we investigated the dependence of Orai2- and Orai3-mediated I_CRAC_ on pH_i_ and compared it to pH_i_ dependence of Orai1. The main results demonstrate that the amplitude of I_CRAC_ mediated by Orai1 and Orai2 strongly depends on pH_i_, whereas the amplitude of I_CRAC_ mediated by Orai3 does not. Furthermore, in the presence of EGTA in the pipette solution, Orai1-mediated I_CRAC_ displays a significant increase in re-activation at negative potentials at acidic pH_i_ (Gavriliouk *et al*., 2017). In contrast, neither Orai2 nor Orai3 exhibits re-activation at any pH_i_. Domain swapping between Orai1 and Orai3 revealed that replacing separately Orai1 N- or C-termini, or intracellular loop 2 with those of Orai3 (Orai1-[Orai3-Nt], Orai1-[Orai3-Ct], and Orai1-[Orai3-L2] chimeras, correspondingly) did not abolish Orai1 dependence on pH_i_. However, replacing both N-terminus and intracellular loop2 in Orai1 with the corresponding parts of Orai3 (Orai1-[Orai3-Nt-L2] construct) rendered Orai1 pH_i_ independent. Conversely, replacing both N-terminus and intracellular loop2 in Orai3 with the corresponding parts of Orai1 (Orai3- [Orai1-Nt-L2] chimera) conferred pH_i_ dependence on Orai3. A significant reduction of I_CRAC_ dependence on pH_i_ seen in H134A Orai1 mutant suggested that in addition to N-terminus and loop 2 of Orai1, some parts of STIM1, most likely within STIM1-CAD, are also involved in I_CRAC_ modulation by intracellular pH.

Although I_CRAC_ recorded in cells co-expressing STIM1 and any of Orai homologues exhibit very similar properties, there are some notable differences in the rate and the extent of FCDI, the inhibition and potentiation by 2-APB and other chemicals, re-activation at negative potentials, sensitivity to reactive-oxygen species, and downstream Ca^2+^ signalling (Lis *et al*., 2007; Bogeski *et al*., 2010; Frischauf *et al*., 2011; Shuttleworth, 2012; Yoast *et al*., 2020; Zhang *et al*., 2020). The results of this investigation add another property that differentiates Orai homologues – dependence on pH_i_. This difference in pH_i_ sensitivity between Orai1 and Orai3 allowed identification of N-terminus and loop 2 of Orai1 as the structures that mediate pH_i_ dependence. Importantly, it is not just specific amino acid sequences in N-terminus and loop 2, but the functional interaction between N-terminus and loop 2 which are important for pH_i_ dependence, as separate replacements of each, N-terminus and loop 2 of Orai1, with the Orai3 counterparts had little effect. Cooperative action of the intracellular domains was previously found to underlie the differences between FCDI and re-activation of Orai1 and Orai3 (Frischauf *et al*., 2011). Attempts to identify which regions of Orai3 make its kinetics distinctly different from that of Orai1 established that all three intracellular domains (N-terminus, loop2 and C-terminus) of Orai1 have to be replaced with those of Orai3 to make kinetics of Orai1 chimera indistinguishable from that of Orai3 (Frischauf *et al*., 2011). In our hands, swapping Orai1 N-terminus and loop2 with those of Orai3 was sufficient to make FCDI of the chimera resemble that of Orai3 and at the same time to remove Orai1 sensitivity to pH_i_, also making it similar to Orai3. Furthermore, replacement of N-terminus and loop2 in Orai3 with those of Orai1, imparted a strong pH_i_ dependence to Orai3, confirming the critical role of N-terminus and loop2 in Orai1-mediated I_CRAC_ dependence on pH_i_.

Both, N-terminus and loop2 of Orai have been shown to interact with STIM1, and the interactions between N-terminus, loop2 and STIM1 are Orai-isoform specific (Fahrner *et al*., 2018). It is possible that isoform specific I_CRAC_ dependence on pH_i_ could be due pH dependence of Orai-STIM1 interactions (Gavriliouk *et al*., 2017). Replacing STIM1 with CAD linked to Orai1 in 1:1 stoichiometry did not abolish pH_i_ dependence of I_CRAC_, suggesting that if pH_i_ has any effect on STIM1-Orai1 interactions, the STIM1 structures involved in these interactions must be localised within CAD. The contribution of STIM1 into Orai1-mediated I_CRAC_ dependence on pH_i_ became more apparent in the experiments using STIM1-independent mutant of Orai1, H134A (Frischauf *et al*., 2017). In the absence of exogenous STIM1, H134A-mediated I_CRAC_ completely lost its potentiation by intracellular alkalinisation, and its inhibition by acidification was significantly attenuated. This reinforces the notion that pH_i_ affects functional coupling of STIM1 and Orai1, which explains the similarities between the effects of intracellular acidification and increased expression of Orai1 relative to STIM1 on I_CRAC_ kinetics and the apparent *P*_o_ curves (Scrimgeour *et al*., 2009; Gavriliouk *et al*., 2017). Employment of H134A mutant also suggested that a smaller, but sizable part of Orai1-mediated I_CRAC_ pH_i_ dependence (∼ 40% inhibition of H134A *vs* ∼ 95% inhibition of WT Orai1 by 60 mM Propionate, EGTA in the pipette) is intrinsic to Orai1. This intrinsic pH_i_ dependence of Orai1 must also be restricted to N-terminus and loop2, but single mutations of protonatable residues in the intracellular domains of Orai1 did not bring any answers thus far.

Histidine 155 in Orai1 loop2, which is conserved in all Orai isoforms, has previously been suggested to be responsible for pH_i_ dependence, particularly the potentiation by alkaline pH_i_, of Orai1-mediated Na^+^ current in the absence of extracellular Ca^2+^ (Tsujikawa *et al*., 2015). In this work, however, it was found that in the presence of 10 mM Ca^2+^ in the bath solution, I_CRAC_ mediated by H155Y Orai1 mutant exhibited pH_i_ dependence virtually identical to that of WT Orai1. Some effects of this mutation on I_CRAC_ kinetics and hence on the apparent *P*_o_, particularly evident in the presence of BAPTA in the pipette, are likely due to the fact that H155 belongs to a stretch of amino acids (153NVHNL157) indispensable for FCDI (Srikanth *et al*., 2010). It is possible, however, that lowering extracellular Ca^2+^ concentration to nanomolar levels affects Orai1-STIM1 interactions to such extent that a single H155Y mutation in loop2, which is critical for pH_i_ dependence, is sufficient to abolish I_CRAC_ potentiation by intracellular alkalinisation, even though the protonation status of His155 may have no bearing on I_CRAC_ amplitude or kinetics.

As mentioned above, both intracellular acidification and high Orai1:STIM1 expression ratio increase the extent of re-activation of Orai1-mediated I_CRAC_ (Scrimgeour *et al*., 2009; Hoover *et al*., 2011; Gavriliouk *et al*., 2017). Furthermore, I_CRAC_ exhibiting strong re-activation due to the high relative expression levels of Orai1 is more sensitive to pH_i_ changes than I_CRAC_ in cells with low Orai1:STIM1 expression ratio (Gavriliouk *et al*., 2017). Experiments with Orai1-CAD revealed that acidic pH_i_ causes closure of Orai1 channels at positive potentials, shifting the apparent *P*_o_ curve to hyperpolarising potentials, which results in more robust re-activation seen during voltage steps to negative potentials. These data also imply that re-activation is not a ‘reversal’ of FCDI, but a separate Ca^2+^ dependent gating process. Presence of re-activation in Orai1 and Orai3-[Orai1-Nt-L2] chimera and its absence in Orai3 and Orai1-[Orai3-Nt-L2] chimera correlated with the presence and absence of pH_i_ dependence, correspondingly. Although the time course of re-activation seen in Orai3-[Orai1-Nt-L2] chimera was two orders of magnitude faster than that of Orai1, it depended on pH_i_ similarly to re-activation of Orai1-mediated I_CRAC_ (Gavriliouk *et al*., 2017). The correlation between pH_i_ dependence and re-activation did not hold for Orai2, however, which exhibited no re-activation but strong pH_i_ dependence.

A potentially physiologically important difference between Orai1, Orai2 and Orai3 is their sensitivity to reactive oxygen species (ROS) (Frisch *et al*., 2019). It has been shown that preincubation of HEK293 cells heterologously expressing STIM1 and either Orai1 or Orai2 in H_2_O_2_ inhibits the development of I_CRAC_. However, I_CRAC_ mediated by Orai3 is insensitive to H_2_O_2_ (Bogeski *et al*., 2010). Ca^2+^, ROS and pH are critical components of tumour microenvironment (Glitsch, 2011; Swietach *et al*., 2014; Pethő *et al*., 2020). Considering that different Orai isoforms have different sensitivity to ROS and pH_i_, it’s quite possible that they make different contributions into the development of a specific tumour microenvironment, and this microenvironment, in turn, determines what roles different Orai isoforms play in cancer development and progression (Frisch *et al*., 2019).

In conclusion, regulation of Orai1 mediated I_CRAC_ amplitude and kinetics by intracellular pH is quite complex and involves several interacting sites on Orai1 and STIM1, including Ca^2+^ binding sites, that have both activating and inactivating actions on I_CRAC_. In contrast, pH_i_ regulates Orai2 amplitude, but not its kinetics, whereas Orai3 does not depend on pH_i_ at all. How this is relevant to CRAC channels function in physiology and pathology remains to be investigated.

## Availability of data and materials

All data and materials used in the current study are available from the corresponding author upon request.

Grigori Rychkov,

Associate Professor,

School of Medicine, University of Adelaide, Adelaide, South Australia 5005, Australia

Telephone: +61-8-81284854;

## Acknowledgements

We would like to thank Dr Lixia Yue, University of Connecticut, USA, for providing H155F mutant of Orai1, Prof Stefan Feske, New York University, NY, USA, for providing pEYFP-C1-Orai1 plasmid and Dr Irene Frischauf and Prof Chris Romanin, University of Linz, Austria, for providing Orai1-[Orai3-L2] and Orai3-[Orai1-Nt-L2] chimeras and H134AOrai1 mutant. We would also like to thank Dr Michael Duffield and Dr Nathan Scrimgeour for conducting preliminary patch clamping experiments.

## Conflict of interests

The authors declare no conflict of interests with the contents of this article.

## Author contributions

GYR designed the experiments; GYR, FHZ, MA and LM conducted the experiments, GYR, SMB and GJB analysed the results and wrote the paper. All authors contributed to manuscript editing.

## Funding

This research was supported by the Australian Research Council Discovery Project 140100259

## Abbreviations

STIM: stromal interaction molecule
CRAC: Ca^2+^ release activated Ca^2+^
2-APB: 2- Aminoethyl diphenylborinate
pH_i_: intracellular pH
SOCE: store-operated Ca^2+^ entry
FCDI: fast Ca^2+^ dependent inactivation
ID_STIM_: STIM1 inactivation domain
CAD: CRAC activation domain
SOAR: STIM1 Orai activating region

## Notes

### Competing Interest Statement

The authors have declared no competing interest.

