## Supporting Information for "Orai1- and Orai2-, but not Orai3-mediated I_CRAC_ is regulated by intracellular pH"

**Supporting Information list:**

**Supplementary Experimental Procedures**
**Supplementary Table**
**Supplementary Figure 1**
**Supplementary Figure 2**
**Supplementary Figure 3**
**Supplementary Figure 4**

**Supplementary Experimental Procedures**

***Generation of Orai1-[Orai3] chimeras***

Chimeras: 1. Orai1-[Orai3-Nt] (Orai1 with N-terminus of Orai3); 2. Orai1-[Orai3-Ct] (Orai1 with C-terminus of Orai3); and 3. Orai1-[Orai3-Nt-L2] (Orai1 with N-terminus and intracellular loop2 of Orai3) were produced by performing PCR reactions using Platinum Taq or Phusion High Fidelity DNA polymerase (Thermo fisher scientific (Life Technologies Australia, Vic, Australia), and sets of primers designed to specifically include the regions of interest in Orai1 and Orai3 and the restriction sites (NheI and EcoRI) according to manufacturer's protocol. Chimera 1, Orai1-[Orai3-Nt], and chimera 2, Orai1-[Orai3-Ct], were produced using Orai1-pAdTrack-CMV and Orai3-pAdTrack-CMV as the DNA templates, while chimera 3- Orai1-[Orai3-Nt-L2] was produced using Orai1-[Orai3-Nt] -pAdTrack-CMV and Orai3-pAdTrack-CMV as templates. The sets of primers and vector templates used are listed in Supplementary Table 1. Specifically, to synthesize Orai1-[Orai3-Nt], set 1 and set 2 primers were used to generate Orai3N fragment with Orai1 overhang at the 3' end and Orai1C fragment with Orai3N overhang at the 5' end, respectively; Orai1-[Orai3-Ct], set 3 and set 4 primers were used to generate Orai1N fragment with Orai3C overhang at the 3' end and Orai3C fragment with Orai1 overhang at the 5' end, respectively; and Orai1-[Orai3-Nt-L2], set 5, set 6 and set 7 primers were used to generate Orai3N-Orai1 fragment with Orai3-Loop2 overhang at the 3'end, Orai3-Loop2 fragment with Orai1 overhang at both the 5' and 3' end and Orai1C fragment with Orai3-Loop2 at the 5' end, respectively in separate PCR reactions. The PCR products (set 1, 2, 3, 4, 5, 6, and 7) were purified using QIA quick gel extraction Kit (Qiagen, Vic, Australia). Additional PCR reactions were performed to combine set 1-2 fragments, set 3-4, and set 5-6-7 fragments to generate chimera 1 (Orai1-[Orai3-Nt]) using set 1 forward and set 2 reverse primers, chimera 2 (Orai1-[Orai3-Ct]) using set 3 forward and set 4 reverse primers, and chimera 3 (Orai1-[Orai3-Nt-L2]) using set 5 forward and set 7 reverse primers, respectively. After PCR products were further gel purified, the vector pCIneo and the fragments of chimeras 1, 2 and 3 were double restrict digested (DRD) with NheI and EcoRI restriction enzymes (NEB, MA, USA) according to manufacturer's guidelines. In order to ligate the chimera inserts into the cut pCIneo vector, the individual DRD chimeras and the DRD-pCIneo vector were gel purified again and treated with T4 DNA ligase (NEB, MA, USUA) in separate reactions, followed by heat inactivation at 65$^{o}$C for 10 min. Ligation products were transformed into subcloning efficient competent DH5α *E. Coli* cells (Thermo fisher Scientific (Life technologies), Vic, Australia), and grown on ampicillin (100 µg/ml) selective plates according to manufacturer's protocol. At least four colonies were picked and grown in LB selective media for plasmid DNA purification using the QIA prep spin miniprep kit (Qiagen, Vic, Australia). All plasmids were sent to Australian Genome Research Facility (AGRF, SA, Australia) for validation by DNA Sanger sequencing.

***Site-directed mutagenesis of Orai1***

EE162/166QQ and H171Y mutations in human Orai1 were generated using forward and reverse primers carrying the specific nucleotide(s) change (Table 1), and pEYFP-C1-Orai1 template (gift from Dr Stefan Feske, New York University, NY, USA) with Phusion High Fidelity DNA polymerase (Thermo fisher scientific (Life Technologies Australia), Vic, Australia) in a PCR reaction according to the manufacturer's instructions. PCR products were digested with Dnp1 (NEB, MA, USA) for 3 hr, transformed into subcloning efficient competent DH5α*E. Coli* cells (Thermo fisher Scientific (Life technologies), Vic, Australia) and grown on kanamycin (30 µg/ml) selective plates according to manufacturer's protocol. At least four colonies were picked and grown in LB selective media for plasmid DNA purification using the QIA prep spin miniprep kit (Qiagen, Vic, Australia). All plasmids were sent to Australian Genome Research Facility (AGRF, SA, Australia) for validation by DNA Sanger sequencing.

**Supplementary Table 1**. Forward and reverse primers for the construction of Orai1-[Orai3-Nt], Orai1-[Orai3-Ct], and Orai1-[Orai3-Nt-L2] chimeras, and for Orai1 mutations (base change is highlighted in **yellow)**.

| **Primers** | **Sequence (5' to 3')** | **Vector Template** |
|---|---|---|
| Set 1-5'NheI-Orai3-Nt3'-FW | CTAGCCGCTAGCACCATGAAGGGCGGCGAGG | Orai3-pAdTrack-CMV |
| Set 1-3'Orai3-Nt-Orai1-Nt5'-RV | TCAAGTAGAGGCGGCGCCAGCTGAGCGCCC | Orai3-pAdTrack-CMV |
| Set 2-5'Orai3-Nt-Orai1-Nt3'-FW | CTGGCGCCGCCTCTACTTGAGCCGCGCCAAG | Orai1-pAdTrack-CMV |
| Set 2-3'Orai1-Ct-stop codon-EcoRI5'-RV | CGGAATTCTCACTAGGCATAGTGGCTGCC | Orai1-pAdTrack-CMV |
| Set 3-5'NheI-Orai1-Nt3'-FW | CTAGCCGCTAGCACCATGCATCCGGAGCCCGCC | Orai1-pAdTrack-CMV |
| Set 3-3'Orai1-Ct-Orai3-Ct5'-RV | TCTTGTGTGCAACCAGTGAGCGGTAGAAGTGG | Orai1-pAdTrack-CMV |
| Set 4-5'Orai1-Ct-Orai3-Ct3'-FW | CTCACTGGTTGCACACAAGACAGACCGCTAC | Orai3-pAdTrack-CMV |
| Set 4-3'Orai3-Ct-stop codon-EcoRI5'-RV | CGGAATTCTCATCACACAGCCTGCAGCTCC | Orai3-pAdTrack-CMV |
| Set 5-5'NheI-Orai3-Nt3'-FW | CTAGCCGCTAGCACCATGAAGGGCGGCGAGG | Orai1-[Orai3-Nt]-pAdTrack-CMV |
| Set 5-3'Orai1-Orai3-L2-5'-RV | GCAGACACGTGGAGATCATGAGCGCAAACAGGTG | Orai1-[Orai3-Nt]-pAdTrack-CMV |
| Set 6-5'Orai1-Orai3-L2-3'-FW | GCTCATGATCTCCACGTGTCTGCTGCCCCAC | Orai3-pAdTrack-CMV |
| Set 6-3'Orai3-L2-Orai1-5'-RV | AGCGTGCCGATGGCAGTGGAGAAGCCC | Orai3-pAdTrack-CMV |
| Set 7-5'Orai3-L2-Orai1-3'-FW | TTCTCCACTGCCATCGGCACGCTGCTCTTC | Orai1-[Orai3-Nt]-pAdTrack-CMV |
| Set 7-3'Orai1-Ct-stop codon-EcoRI5'-RV | CGGAATTCTCACTAGGCATAGTGGCTGCC | Orai1-[Orai3-Nt]-pAdTrack-CMV |
| Orai1-EE162/166QQ-FW | CTCAACTCGGTCAAG**C**AGTCCCCCAT**C**AGCGCATGCACCGCC | pEYFP-C1-Orai1 |
| Orai1-EE162/166QQ-RV | GGCGGTGCATGCGCT**G**ATGGGGGACT**G**CTTGACCGAGTTGAG | pEYFP-C1-Orai1 |
| Orai1-H171Y-FW | ATGCACCGC**T**ACATCGAGCTGGCCTGG | pEYFP-C1-Orai1 |
| Orai1-H171Y-RV | CTCGATGT**A**GCGGTGCATGCGCTCATGG | pEYFP-C1-Orai1 |

### Supplementary Figure 1

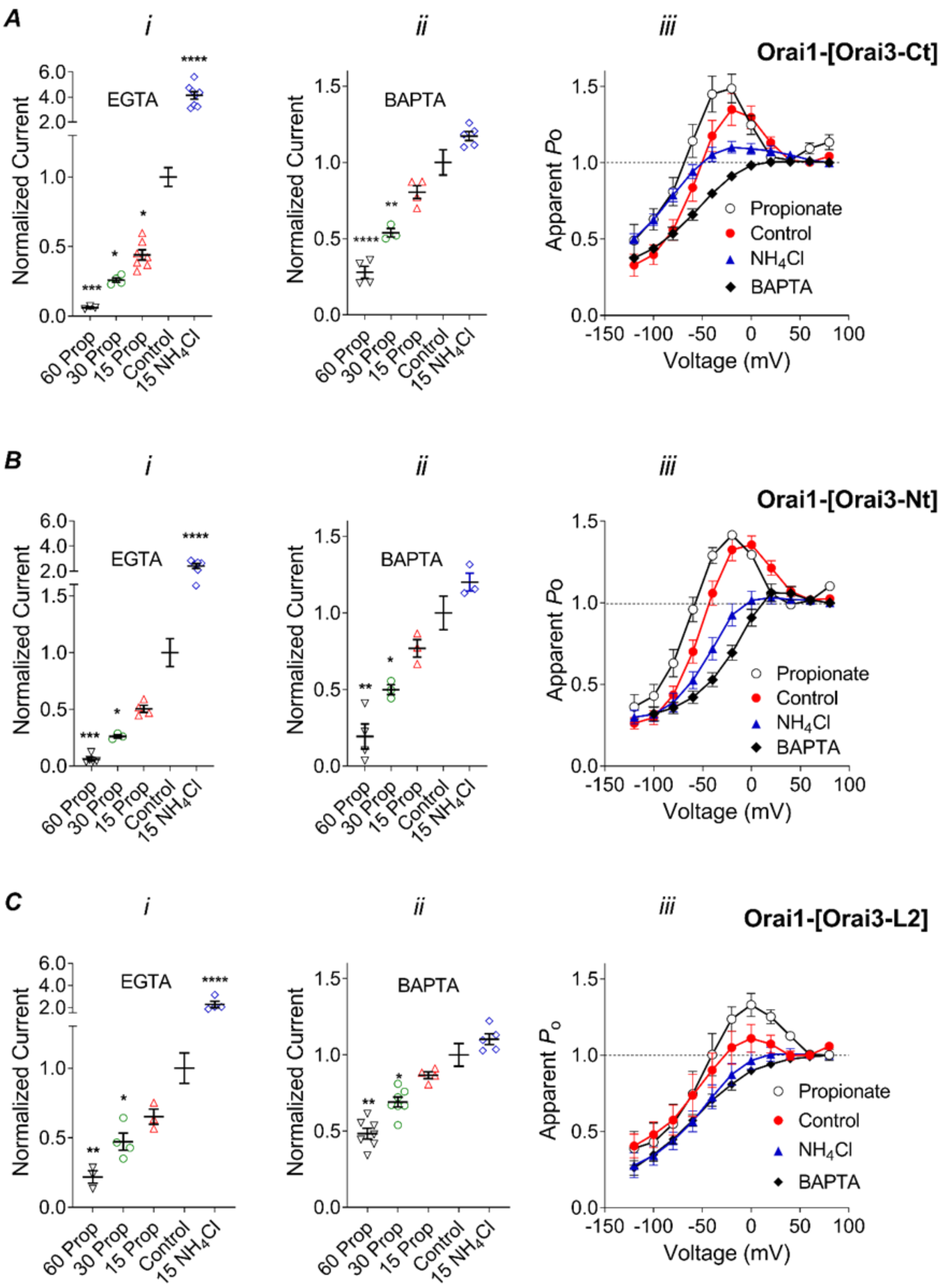


**Figure S1. pH$_i$ dependence of Orai1-[Orai3-Ct] (A), Orai1-[Orai3-Nt] (B), and Orai1-[Orai3-L2] (C) chimeras.** ***i*, *ii*.** Average normalised current amplitude in the presence of Na Propionate or $NH_4Cl$ in the bath and either EGTA (*i*) or BAPTA (*ii*) in the pipette solution. ***iii*,** Apparent $P_o$ curves obtained from current recordings in the control bath solution, in the presence of 15 mM Na Propionate, or 15 mM $NH_4Cl$ in the bath and EGTA in the pipette solution, and in the control bath solution and BAPTA in the pipette solution. Data was analysed using One-way ANOVA with multiple comparisons indicated within the panels.

**Supplementary Figure 2**

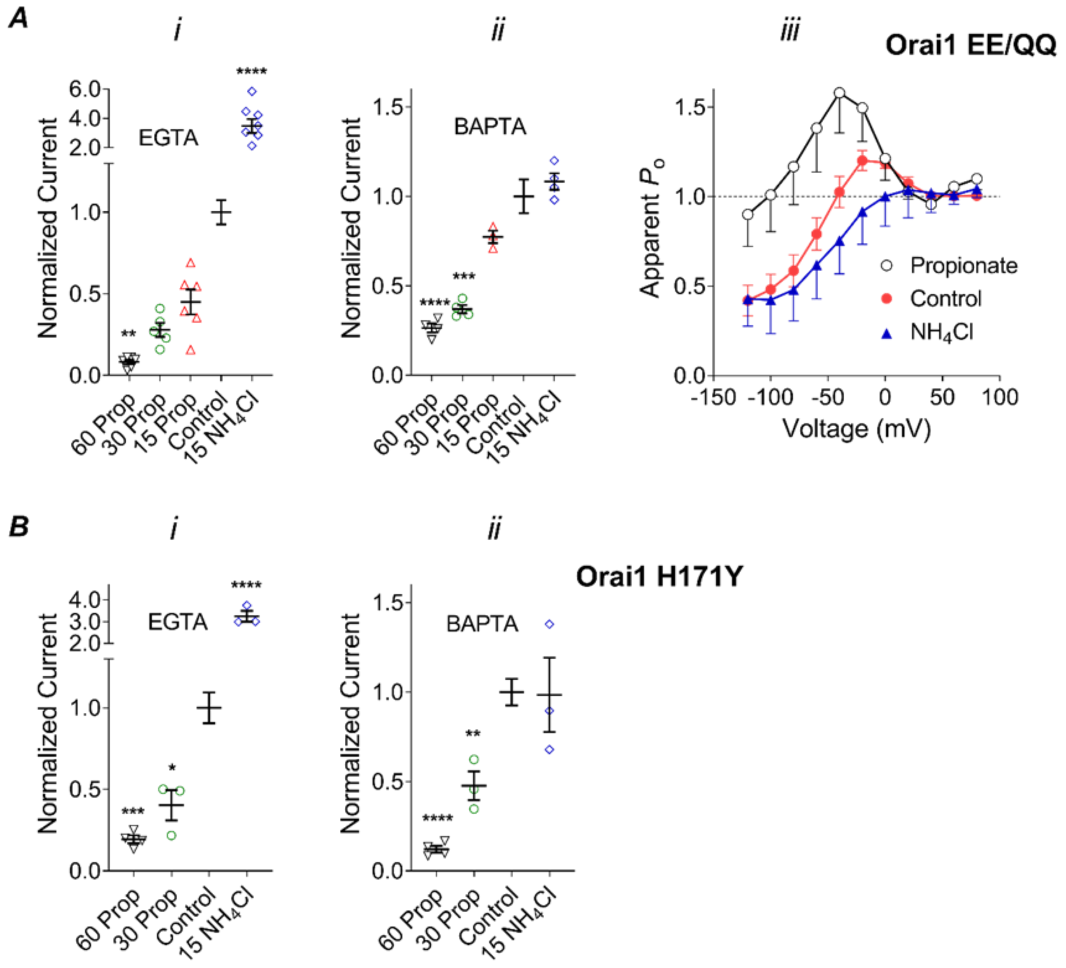


**Figure S2. pHi dependence of 162E,166E/QQ (A) and H171Y (B) Orai1 mutants**. ***i***, ***ii***. Average normalised amplitude of $I_{CRAC}$ in the presence of Na Propionate or $NH_4Cl$ in the bath and either EGTA (*i*) or BAPTA (*ii*) in the pipette solution. Data was analysed using One-way ANOVA with multiple comparisons indicated within the panels. ***iii***. Apparent $P_o$ curves of 162E,166E/QQ-Orai1 recorded in the control bath solution, in the presence of 15 mM Na Propionate or 15 mM $NH_4Cl$ in the bath and EGTA in the pipette solution, as indicated on the panel.

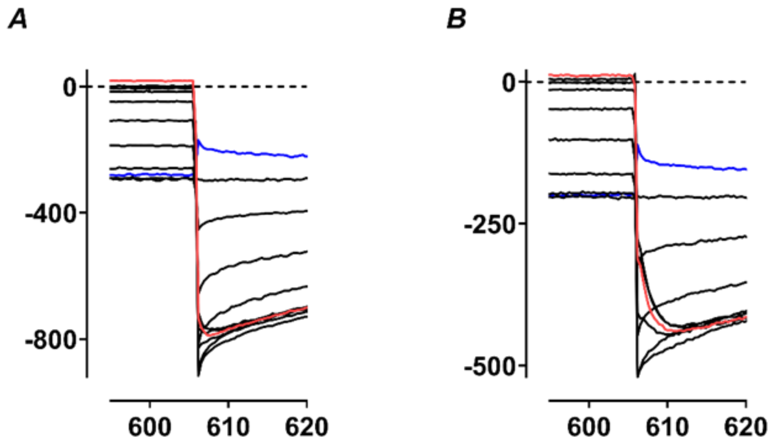


**Figure S3. Re-activation kinetics of Orai3-[Orai1-Nt-L2] $I_{CRAC}$.** Representative Orai3-[Orai1-Nt-L2] $I_{CRAC}$ traces used to construct apparent $P_o$ curves in the control bath solution (A) and bath solution with 15 mM propionate (B) at expanded time scale.

**Supplementary Figure 4**

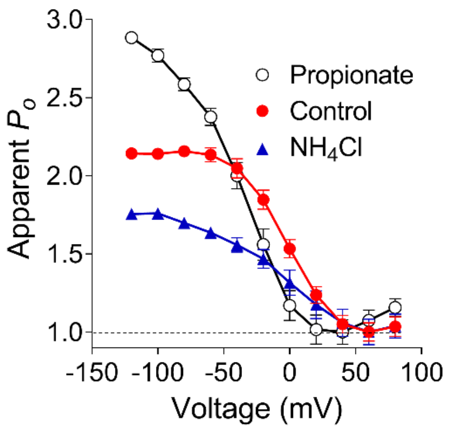


**Figure S4. Apparent $P_o$ curves of Orai1-CAD construct**. These are the same data as shown in Figure 6 but normalised to the amplitude of the tail current after a pulse to 60 mV. This is shown to enable a direct comparison with $P_o$ curves of all other constructs in this study.
